# Maternal RhD heterozygous genotype is associated with male biased secondary sex ratio

**DOI:** 10.1101/543629

**Authors:** Šárka Kaňková, Jaroslav Flegr, Jan Toman, Pavel Calda

**Author notes:** Corresponding author, Division of Biology, Faculty of Science, Charles University, Prague, Viničná 7, 128 44, Czech Republic. **Declarations of interest** None.

## Abstract

The results of previous studies overwhelmingly suggest that RhD positive heterozygotes express better health status than Rh positive homozygous, especially in RhD negative subjects. This also applies to pregnant women. According to the Trivers-Willard hypothesis, women in better physical condition should have a male-skewed sex ratio. The aim of the present study was to test the hypothesis that RhD positive heterozygous mothers give birth to more sons than daughters. In the present cross-sectional study, we analysed data from 5,655 women who have given birth in the General University Hospital in Prague, Czech Republic between 2008-2012. Clinical records comprised maternal weight before pregnancy, number of previous deliveries, sex of the newborn, maternal RhD phenotype, and RhD phenotype of the newborn. Secondary sex ratio was significantly higher (P=0.028) in RhD positive mothers who had RhD negative newborns, i.e. in heterozygotes (SR=1.23), than in RhD positive mothers who had RhD positive newborns, i.e. in a mixed population of heterozygotes and homozygotes (SR=1.00), especially in primiparous women (P=0.013; SR=1.37 and 0.99 resp.). In line with the Trivers-Willard effect, RhD maternal heterozygous genotype is associated with male biased secondary sex ratio. The results supported the hypothesis that RhD polymorphism may be maintained due to heterozygote health advantages.

## 1. Introduction

The RhD protein, a product of the *RHD* gene, is a major component of the Rh blood group system. It carries the strongest blood group immunogen, which is the D antigen. Considering its role, the molecular structure of the RhD protein suggests it is a part of an ion pump present in the red blood cell membrane. The whole complex probably serves for the transport of NH_3_ or CO_2_ molecules across the erythrocyte cell membrane (Flegel, 2011; Kustu and Inwood, 2006). However, its physiological role is still unclear. Several possible options have been discussed, e.g. in Flegr et al. (2015).

Almost 85 % of Europeans express an RhD positive phenotype – the RhD protein is present in their erythrocyte cell membranes. However, the RhD antigen is absent in a considerable portion of the European population (RhD negative subjects) due to the *RHD* deletion (Wagner and Flegel, 2000). RhD polymorphism is comparably high in numerous other human populations (Golassa et al., 2017; Mourant, 1976), but its existence is an evolutionary enigma. Theoretically, populations should be RhD monomorphic. This is because of a strong selection against RhD positive children born to RhD negative mothers due to hemolytic disease of the newborn (Bowman, 1997; Filbey et al., 1995). Before the introduction of prophylactic treatment, this disorder, which can cause serious illness, brain damage, or even death of the fetus or newborn of multiparous women, was one of the leading causes of newborn mortality in highly RhD polymorphic populations. Therefore, the representatives of the minor phenotype had lower fitness before the advent of modern medicine. This includes either RhD negative women in a mostly RhD positive population or RhD positive men in a mostly RhD negative population.

Erythrocytes of RhD- and RhD+ homozygotes differ in the molecular complexes present on their cell membranes, and most likely in their biological activities as well (Le Van Kim et al., 2006; Wagner and Flegel, 2000). Moreover, a difference in the erythrocytes of RhD positive homozygotes and heterozygotes was also observed. About 33,560 D antigen sites were detected on the surface of RhD positive homozygous erythrocytes, whereas the surface of RhD heterozygous erythrocytes contain about 17,720 D antigen sites (McGann et al., 2012). It is thus possible that the vulnerability of RhD positive homozygotes, RhD positive heterozygotes and, especially, RhD negative homozygotes to various irregular conditions, including various diseases, may differ dramatically. RhD heterozygotes may link the advantages of both RhD negative and positive homozygotes and the RhD polymorphism in human populations thus could be maintained by selection in favour of heterozygotes (Feldman et al., 1969; Fincham, 1972).

The correlation of particular RhD genotypes with specific health conditions has been supported by several studies (see, e.g. Flegr et al., 2008, 2009, 2010, 2012, 2013; Kankova et al., 2010; Novotna et al., 2008). One interesting theme is that RhD negativity and heterozygosis often affect the individual’s health in opposite directions. Recently, two independent studies showed that RhD phenotype strongly affects the incidence and prevalence of many disorders. RhD negative subjects reported having more frequent allergic, digestive, heart, haematological, immunity, mental health, and neurological problems, as well as a higher incidence of some infectious diseases. This was corroborated by the reported frequency of their visits to medical specialists, usage of prescribed drugs, headaches, and general tiredness (Flegr et al., 2015). The results showed a complex picture. Certain significantly elevated medical problems were specific to RhD negative individuals, others to RhD positive persons, most of them in a sex-specific way. However, taken together, the RhD negative subjects had more serious health problems than the RhD positive subjects in all six variables, significantly differing according to RhD of 22 variables analysed. The difference in RhD phenotype was obviously caused by the underlying RhD genotype, but RhD positive homozygotes and heterozygotes were not separated in this study. Some of the results indicate that RhD negative phenotype may confer increased immunity to infections of viral origin. In the light of these results and considering the long-term persistence of RhD polymorphism, however, it seems more probable that it could be RhD positive heterozygotes who are selectively advantageous in human populations. RhD polymorphism thus could be sustained in populations by negative frequency-dependent selection, namely by its specific form – the selection in favour of heterozygotes (heterozygous advantage).

A recent ecological regression study performed on a set of 65 countries for which the RhD genotype frequencies data were available, showed the strongest evidence yet for the heterozygote advantage hypothesis (Flegr, 2016). The results showed that both the frequencies of RhD negative homozygotes and RhD positive heterozygotes (whose frequency was calculated from data on the frequency of homozygotes using Hardy-Weinberg equation) correlated with specific disease burdens in particular countries. In general, the burdens (both in the sense of Disability Adjusted Life Year, see WHO, 2008, and disease mortality rates) were higher in RhD negative homozygotes. Moreover, the observed correlations mostly lead in opposite directions in RhD negative homozygotes and RhD positive heterozygotes. The general pattern showed that the countries with a high frequency of Rhesus negative homozygotes had a lower burden associated with congenital anomalies and neuropsychiatric conditions. Additionally, such countries had a higher burden of cardiovascular diseases and, especially, of malignant neoplasm. This strongly supports the hypothesis that RhD polymorphism could be sustained in populations by selection in favour of heterozygotes at the expense of alternative hypotheses that consider its current distribution to be a consequence of founder effects, genetic disequilibrium, and/or gene flow attenuated by viability selection, partial reproductive incompatibility, and reproductive compensation (see, e.g. Feldman et al., 1969; Nei et al., 1981).

Several studies directly supporting RhD heterozygous advantage have also been published. Flegr et al. (2008) and Novotna et al. (2008) showed that healthy RhD negative homozygotes exhibit faster reactions than RhD positive subjects. This reaction time, however, strikingly changes when subjects were infected by *Toxoplasma gondii,* a common parasite whose chronic prevalence in various countries ranges between ten and ninety percent. *Toxoplasma* positive subjects showed prolonged reaction times. This impairment was most severe in RhD negative homozygotes, whereas RhD positive individuals, and especially RhD positive heterozygotes, were considerably protected from this effect. In consequence, the performance of *Toxoplasma* positive subjects was best in RhD positive heterozygotes followed by RhD positive homozygotes and RhD negative homozygotes. The same data also showed that RhD positivity (most probably RhD heterozygosity) modulates certain effects of smoking, fatigue, and ageing (Flegr et al., 2012). For example, the positive effect of age on performance and intelligence was stronger in RhD positive subjects, whereas the effect of smoking on the number of viral and bacterial diseases was about three times higher in RhD negative subjects. In a different study, *Toxoplasma* positive RhD negative subjects expressed lower performance in weight-holding and hand-grip tests in comparison with both RhD negative and positive *Toxoplasma* negative subjects (Flegr et al., 2018). It is clear that faster reaction times or greater stamina conferred a selective advantage in the past, and that they still do in the present. It was, for example, demonstrated that the probability of being involved in a traffic accident is elevated more than twice (in the case of chronic infection) or even five times (in the case of recent infection) in *Toxoplasma* positive RhD negative homozygotes (Flegr et al., 2009). This could lead to the spread of the RhD negative allele in a similar manner to the recessive *HBB* allele for sickle cell disease. *HBB* recessive homozygotes have severely impaired viability. In a heterozygous condition, the *HBB* allele does not impair viability in most situations but confers an increased resistance to malaria. The *HBB* allele thus spread in tropical areas with *Plasmodium falciparum* by the means of frequency-dependent selection, or its specific form – heterozygous advantage – until a disproportionate fraction of nonviable recessive homozygotes are born and its frequency in a population is stabilized (Allison, 1954).

However, phenotypic correlates of RhD phenotype may be more complex. A study performed on 502 soldiers surprisingly showed that *Toxoplasma* positive RhD positive subjects express lower, while *Toxoplasma* positive RhD negative subjects express higher, verbal and nonverbal intelligence than their *Toxoplasma* negative peers (Flegr et al., 2013). This may be the consequence of psychological differences between *Toxoplasma* positive and *Toxoplasma* negative subjects (a lower total N-70 score of potentially pathognomic factors, anxiety, depression, phobia, hysteria, vegetative lability, hypochondria, psychasteny, and neuroticism, but see Flegr et al., 2010, for somewhat contradictory results) that were observed in this study and were more prominent among RhD negative subjects. However, these psychological differences may result from the increased tendency of *Toxoplasma* positive military personnel to mask any negative properties. Moreover, there were signs of higher verbal and nonverbal intelligence even in *Toxoplasma* negative RhD negative subjects, which points to a possible direct effect of RhD phenotype. Regardless, the protective role of RhD positive phenotype against the effects of toxoplasmosis was documented even in this study. The strongest effect of Rh phenotype was reported in a study on the influence of *Toxoplasma* on weight gain in pregnancy. In the study of Kankova et al. (2010), RhD negative women with latent toxoplasmosis gained nearly two times more weight in 16^th^ week of pregnancy (N = 27) than RhD negative *Toxoplasma* negative women (N = 139) or the RhD positive women of any infection status (N = 813). The difference of about 1600 g remained approximately constant until delivery.

The secondary sex ratio (sex ratio at birth) in humans is around 1.06 in most populations (Davis et al., 1998). Within the population, the sex ratio may be influenced by many factors, such as paternal hormones (James, 1996, 2010, 2015; James and Grech, 2018), immunosuppression (James, 1996), and several important pathologies (e.g. hepatitis, James, 2010, or toxoplasmosis, Kankova et al., 2007). As the Rh phenotype modulate the effects of many detrimental factors on human performance and physiology, we decided to examine its potential effects on sex ratio at birth. The specific aim of the present study was to analyse the association between the sex of newborns and RhD phenotype of both the newborns and mothers. Our working hypothesis was to expect a higher secondary sex ratio in RhD heterozygous women. The generally accepted Trivers-Willard hypothesis (Trivers and Willard, 1973) suggests that females, including women in “good condition”, e.g. women with a good health status, tend to give birth to more sons than daughters. The reason is that mothers in good condition may invest disproportionately more time, energy, and resources into their sons, which may, in turn, reach a better condition and leave more offspring in polygynous (or serially monogamous) species where the biological fitness of males, but not so much females, strongly depends on their health status. According to the studies, heterozygous RhD positive mothers probably express better health status than Rh positive homozygous mothers and especially than the RhD negative mothers. This can result in a higher sex ratio in RhD positive than in RhD negative mothers and an even higher sex ratio in RhD positive mothers who give birth to RhD negative children, i.e., in RhD heterozygous mothers. We expect this effect to be more prominent in primiparous women on the basis of already published results (Christiansen et al., 2004; Maraz et al., 1973; Nielsen et al., 2008).

## 2. Material and Methods

### 2.1. Subjects

The study was designed as a cross-sectional study. The main data set covered women who have given birth in the General University Hospital in Prague, Czech Republic between 2008-2012. Clinical records comprised maternal weight before pregnancy, number of previous deliveries (primiparae/multiparae), sex of the newborn, maternal RhD phenotype (positive/negative), and RhD phenotype of the newborn (positive/negative). The women that gave birth to twins were excluded from the analyses. During the whole study, we worked with an anonymized data set.

### 2.2. Statistics

The program Statistica 10.0 was used for all statistical testing. The association between the sex of the newborn and the RhD phenotype of the newborn was analysed using the Chi-Square test, initially for all women, and then separately for RhD positive and RhD negative women. Furthermore, the analysis was conducted separately for both primiparous and multiparous women. The effects of maternal weight as a continuous predictor and the RhD phenotype of newborn (positive/negative) as a categorical predictor of newborn sex, were evaluated by the generalized linear model (GLZ) separately for both RhD positive and RhD negative women. Binomial distribution and logit link function, as recommended by Wilson and Hardy (2002), were used for the construction of the model. For some women, the variable “maternal weight before pregnancy” was not available, and therefore the number of women varied between analyses. Sex ratio (SR) in this article is expressed as the ratio of male to female.

### 2.3. Ethical approval

The project was approved by IRB Faculty of Science, Charles University (No. 2018/19) (Etická komise pro práci s lidmi a lidským materiálem Přírodovědecké fakulty Univerzity Karlovy).

## 3. Results

The total data set contained the records of 5,655 women. The relationships between the sex of newborns and the RhD phenotype of the newborns analysed using the Chi-Square test showed only a non-significantly male biased proportion of the newborns in the group of RhD negative newborns (N= 5,655, P=0.088, χ^2^=2.91). Among 1,356 RhD negative newborns, 717 were boys (SR=1.12), while among 4,299 RhD positive newborns, only 2,151 were boys (SR=1.01). However, the situation differed in the RhD positive and RhD negative subpopulations of women. In women with the RhD positive phenotype, who included Rh positive homozygotes and heterozygotes, the relationships between the sex of newborn and the RhD phenotype of the newborn were significant (N=3,406, P=0.028, χ^2^=4.84). Among 422 RhD negative newborns, the offspring of the RhD positive heterozygote mothers, 235 were boys (SR=1.26), while among 2,984 RhD positive newborns, the offspring of either RhD positive heterozygote or RhD positive homozygote mothers, 1493 were boys (SR=0.99). In women with the RhD negative phenotype, the relationships between the sex of the newborns and the RhD phenotype of the newborns did not exist (N=2,229, P=0.660, χ^2^=0.19). Among 929 RhD negative newborns, 479 were boys (SR=1.06), while among 1,300 RhD positive newborns, 658 were boys (SR=1.02).

The same analyses were conducted separately for both primiparous and multiparous women. In primiparous women, the relationship between the sex of the newborn and the RhD phenotype of the newborn analysed using the Chi-Square test showed a significantly male biased proportion of the newborns in the group of RhD negative newborns (N= 3,418, P=0.013, χ^2^=6.21). Among 844 RhD negative newborns, 466 were boys (SR=1.23) and among 2,574 RhD positive newborns, 1,294 were boys (SR=1.01). In women with the RhD positive phenotype, the relationship between the sex of the newborn and the RhD phenotype of the newborn was also significant (N=2,047, P=0.013, χ^2^=6.18). Among 258 RhD negative newborns, 150 were boys (SR=1.37) and among 1,789 RhD positive newborns, 892 were boys (SR=0.99). In primiparous women with RhD negative phenotype, the relationship between the sex of the newborns and the RhD phenotype of the newborns did not exist (N=1,360; P=0.335; χ^2^=0.93). Among 583 RhD negative newborns, 314 were boys (SR=1.17) and among 777 RhD positive newborns, 398 were boys (SR=1.02). In multiparous women, both RhD negative (P=0.559) and RhD positive (P=0.690), no significant results were observed.

In the next part of the study, we analysed the influence of the RhD phenotype of newborns (positive/negative) on the sex of newborns using a generalized linear model (GLZ). At first, this test was conducted only for RhD positive women, while the variables “maternal weight” and “maternal age” were used as the continuous covariates. However, the maternal weight (P=0.998) and the maternal age (P=0.484) did not make a significant contribution to the equation. Finally, the reduced statistical model was conducted without them. The results of GLZ (P values) were similar to the results of the Chi-Square test mentioned above. Again, no significant effect of the RhD phenotype of newborns on the sex of newborns was observed in the group of Rh negative women (data not shown).

## 4. Discussion

Our results showed that the sex ratio at birth was significantly higher (male skewed) in RhD positive mothers who had RhD negative newborns (SR=1.23) than in RhD positive mothers who had RhD positive newborns (SR=1.00). This effect was stronger in primiparous women (SR=1.37 and 0.99 resp.). In our study, we had only data regarding RhD phenotypes. It can be deduced, however, that the RhD negative mothers were undoubtedly homozygotes (dd) while the RhD positive mothers could have either RhD positive heterozygous genotype (Dd) or RhD positive homozygous genotype (DD). This uncertainty applies to the group of RhD positive mothers who had RhD positive newborns. On the other hand, the RhD positive mothers who had RhD negative newborns necessarily had RhD positive heterozygous genotype. Therefore, the main observed result, i.e., higher secondary sex ratio in RhD negative newborns of RhD positive mothers, confirms our *a priori* hypothesis that RhD maternal heterozygous genotype is associated with male biased secondary sex ratio.

Recent results support the hypothesis that the Rhesus factor polymorphism is maintained in human populations due to a higher resistance or tolerance of heterozygotes to specific diseases (see Introduction). The hypothesis was repeatedly supported by empirical data showing that RhD positivity, and especially RhD heterozygosity, protects people against certain negative effects of toxoplasmosis (Flegr et al., 2008, 2018; Kankova et al., 2010; Novotna et al., 2008). In line with the Trivers-Willard effect (Trivers and Willard, 1973), our working hypothesis was to expect RhD heterozygous mothers, i.e., mothers with supposedly higher health status, to exhibit a higher secondary sex ratio. The results of our study therefore supported our working hypothesis and further supported the hypothesis that RhD polymorphism may be maintained due to heterozygote health advantages.

As expected on the basis of already published data (Maraz et al., 1973), the results of our analyses were not significant in multiparous women. This could have been caused by a broader spectrum of factors that influence the secondary sex ratio in multiparous women in comparison with primiparous women. These factors are, for example, sex (Christiansen et al., 2004; Gualtieri et al., 1984; Renkonen et al., 1962) and RhD (Renkonen and Seppala, 1962) of previous siblings, the existence or absence of previous miscarriages (Christiansen et al., 2004; Nielsen et al., 2008), and immunosuppression (James, 1996), especially the reaction against y-antigens (Nielsen et al., 2008, 2009). Due to these sources of latent variability, the effect of RhD phenotype (resp. genotype) on sex ratio at birth cannot be detected in the multiparous women, even if it existed. It must be emphasized, however, that the broader spectrum of these confounding factors cannot be the only reason for the difference in the effect of RhD on primiparous and multiparous women. Confounding factors may increase the variability of the focal variable (the probability of a birth of a son in this study), and by this, they can increase the P value of tests. However, they cannot influence the size of the observed effect. For this reason, our results suggest that effects sizes in primiparous mothers are much higher than in multiparous women, the difference in the effect of RhD on primiparous and multiparous women cannot be explained solely by the aforementioned confounding factors.

Only RhD phenotype, not RhD genotype, of newborns and mothers is examined and recorded in the current clinical praxis. The genotype of RhD negative mothers is dd and the genotype of one third of RhD positive heterozygote mothers (Dd) can deduced from the RhD phenotype of their children. Therefore, we can compute SRB (sex ratio at birth) in these two subpopulations, but not in the subpopulation of RhD positive women with RhD positive children, which represent a mixture of RhD positive homozygotes and heterozygotes. The secondary sex ratio of RhD positive newborns of RhD positive mothers was 0.99. We can expect that the secondary sex ratio in RhD positive newborns of RhD positive mothers with RhD heterozygous genotype (part of RhD positive newborns of RhD positive mothers) is comparable with the secondary sex ratio (SR = 1.37) of RhD negative newborns of RhD positive mothers (with RhD heterozygous genotype). This would then suggest (in the case that the Hardy-Weinberg equilibrium applies and the frequency of RhD negative homozygotes is about 16 % in Czech population, see Flegr, 2016) that the secondary sex ratio in newborns of RhD positive mothers with a RhD positive homozygous genotype is female biased (SR = 0.74) to make the total sex ratio 0.99 for the mixture of RhD positive heterozygotes (SR = 1.37) and RhD positive homozygotes.

It is also worth mentioning that the observed frequency of the RhD negative allele (about 40 % of the standing polymorphism) conspicuously well approaches the proportion of the recessive allele that ensures the production of the relatively highest proportion of heterozygotes (i.e., the most fit individuals; in this case 48 %) and dominant homozygotes (i.e., the individuals with mediocre fitness; 36 %) at the expense of recessive homozygotes (i.e., the least fit individuals; 16 %). The highest possible stable proportion of the most fit heterozygotes in the population is 50 %. However, both dominant homozygotes and the least fit recessive homozygotes would constitute 25 % of the population in this scenario. Gradual change in the proportion of both alleles in favour of dominant one would be expected in this case. The observed deflection in the proportion of RhD alleles thus further supports the hypothesis of the RhD heterozygote advantage and the relatively lowest fitness value of the recessive RhD allele.

Possible alternative scenarios consistent with the observed data suggest it is possible that the observed phenomenon of biased sex ratio in RhD negative children of RhD positive mothers is associated with the child’s RhD genotype. It cannot be excluded that RhD negative embryos are rejected by maternal body less often, which would point to the ability of the RhD allele or some closely linked allele to manipulate the mother’s body or a general preference and selective advantage of RhD negative homozygotes. However, this hypothesis does not seem very probable because the sex ratio of RhD negative mothers was not significantly biased.

Another option is based on the observation that adverse effects of RhD negativity on health probably manifest at an older age (Flegr et al., 2015). This is corroborated by our results (unpublished data), which show that better reaction times in RhD negative individuals are specific to younger age. This points to a possible conditional selective advantage of RhD negative homozygotes. Under this scenario, RhD negativity can be beneficial at a younger age, or at least exhibit its adverse effects on health or psychomotor performance exclusively or more prominently at an older age. Negative effects that manifest only at an older age could hide away into the “selection shadow” (Fisher, 1930; Medawar, 1946). It was supported that mutations that negatively affect fitness at an older age are under a much weaker selective pressure than mutations that affect the fitness of young individuals (Gavrilova et al., 1998). It is also possible that the adverse effects of RhD negative phenotype would be individually reflected during ageing and thus masked by an adaptive behaviour (see, e.g. Flegr et al., 2009, 2013; Novotna et al., 2008). It is even possible that the RhD negative allele represents one of the alleles with antagonistic effects on fitness dependent on age (see, e.g. Pedersen, 1995). Its presence may easily result in a positive feedback loop of selection on fast life strategy and maximum performance at a young age (Nettle, 2010; Promislow and Harvey, 1990). In such a situation, it might be more advantageous to have more RhD negative sons than RhD negative daughters for mothers with RhD positive heterozygous genotypes. Such sons would have a better chance of reproductive success at a young age. In contrast, at least mediocre longevity would be always advantageous for daughters. This presents a special case of the generalized Trivers-Willard hypothesis (Kanazawa, 2005). Note, however, that this option does not exclude advantages of preferentially producing sons by RhD heterozygous mothers in good condition postulated by the classical Trivers-Willard hypothesis (Trivers and Willard, 1973). The benefit of producing RhD negative sons because of their young age advantage therefore might be masked in RhD negative mothers by their own advantage in producing daughters, which results in the pattern we observed in our study.

The important limitation of our study is that we knew only maternal and child RhD phenotypes and not genotypes. We were not able to count maternal RhD genotypes for all data, but only for the subpopulation of 2,651 women (RhD negative women and RhD positive women with RhD negative newborns). It will be necessary to genotype the RhD positive mothers in future studies to confirm or reject our predictions that the SR of RhD positive heterozygotes will be male skewed and the SR of RhD positive homozygotes will be female skewed. The second limitation was a relatively small sample size of sub-analyses evaluated across the different RhD phenotype categories for both primiparous and multiparous women. However, the small number of subjects may lead only to a false negative, not a false positive result of a study. Another, more general, problem was the absence of information on many factors that could have influenced sex ratio at birth (e.g. maternal socio-economic status, health, sex of previous child). Existence of known and unknown confounding factors may result in the failure of a statistical test to detect the existent effect, not in the detection of nonexisting effect (Flegr and Horacek, 2017). However, researchers should focus on these factors in future studies on the role of RhD polymorphism in the origin of skewed human sex ratios.

### 4.1. Conclusions

In our relatively large data set, the maternal RhD heterozygosity was associated with a male biased secondary sex ratio. The most parsimonious explanation of the observed pattern suggests the preference of male offspring by mothers in good condition according to the Trivers-Willard hypothesis. This hypothesis proposes that mothers who exhibit a good physiological condition and/or social status give birth to a higher proportion of sons, while those in a worse situation, e.g. those with worse health, give birth to more daughters. If the explanation is correct, then our data provides new indirect support for the heterozygous advantage hypothesis of sustaining RhD polymorphism in human populations. Moreover, the possibility that the RhD homozygotes, both RhD negative and RhD positive, have worse health than RhD heterozygotes could have serious clinical implications and should be examined in detail in future studies.

## 5. Data Availability

The data associated with this research are available at https://figshare.com/articles/Maternal_RhD_heterozygous_genotype_is_associated_with_mal_e_biased_secondary_sex_ratio/7687544

## Acknowledgment

This work was supported by Czech Science Foundation (grant No. 18-13692S); Charles University (Research Centre program No. 204056); and the Ministry of Health of the Czech Republic (grant RVO-VFN64165). The funding sources were not involved in study design, in the collection, analysis and interpretation of data, in the writing of the report and in the decision to submit the article for publication.

